# Pangenome Analysis of Pyrus Reveals Potential Role of Transcription Factors in Recent Adaptation to Arid Environments

**DOI:** 10.64898/2026.01.20.700718

**Authors:** June Labbancz, Amit Dhingra

## Abstract

Pears (genus *Pyrus*) are among the most extensively cultivated tree fruits with a wide-reaching economic impact. Despite this, the genetic basis of most pear traits of interest, including abiotic stress tolerance, tree architecture, precocity, parthenocarpy, disease resistance, and fruit ripening, remains poorly understood. Although extensive efforts have been made to identify quantitative trait loci (QTLs) that explain the genetic basis of pear traits, many are poorly transferable, limiting their utility for informing genetic improvement or management of pears across most genetic backgrounds. To provide a whole-genome context and enable the exploration of functional variation in *Pyrus*, we developed a pangenome graph using 31 accessions representing 23 *Pyrus* species from the National Clonal Germplasm Repository. Whole-genome sequencing was performed solely with Oxford Nanopore, generating highly contiguous assemblies for pangenome construction, demonstrating the viability of a single-platform approach to pangenomic analysis in Viridiplantae. Exploration of the pangenome graph reveals genes present in some lineages, with potential functional implications. A group of arid-adapted *Pyrus* species exhibits signs of selective sweeps in regions associated with transcription factors, likely impacting abiotic stress tolerance. With the development of this pangenomic resource, resequencing analysis in *Pyrus* is now possible without the limitations imposed by single-reference genome assemblies.

## Introduction

Divergence within *Pyrus* began 25 to 33 million years ago, with the separation of the Occidental and Oriental clades occurring 6 to 19 million years ago, and 8 to 24 million years ago, respectively (Korotkova et al., 2018). *Pyrus* species have diversified in their distribution across much of the mid-latitudes of Eurasia, primarily inhabiting mesophytic forests and xerophytic open woodlands (Korotkova et al., 2018). *Pyrus* phylogenetics is frustrated by the ease with which interspecific hybridization occurs, their overlapping ranges, and apparent morphological similarities; this highly reticulate evolution makes direct linear relationships difficult to resolve (Zheng et al., 2014a; Jiang et al., 2016; Jin et al., 2024a). *Pyrus* has been divided into varying numbers of groups, but the most enduring has been the division into “Occidental” and “Oriental” groups, a division consistently validated by population genetic surveys (Liu et al., 2015; Volk and Cornille, 2019; Montanari et al., 2020; Jin et al., 2024b). Consequently, it has been proposed, with morphological and phylogenomic support, that this division be elevated to the level of subgenera, *Pyrus* subg. *Pyrus* and *Pyrus* subg. *Pashia* (Jin et al., 2024b). *Pyrus regelii* appears to cluster separately from both groupings but has generally been grouped into *Pyrus* subg. *Pyrus* (Zheng et al., 2014b; Quinet and Wesel, 2019; Montanari et al., 2020). The origin of the domesticated European pear (*P. communis*) remains poorly understood beyond its origin in species of *Pyrus* subg. *Pyrus*. In modern times, European pear is grown primarily in temperate to semi-arid regions of Europe, North America, and South America (Quinet and Wesel, 2019). Commercially cultivated Asian pears (*P. pyrifolia, P. ussuriensis, P. bretschneideri*), domesticated from species in *Pyrus* subg. *Pashia,* are grown primarily in East Asia, from humid subtropical to semi-arid climates (Quinet and Wesel, 2019). Despite the genetically distinct origins of the European and Asian pear and their independent domestication events, both are cultivated in temperate climates with similar cultivation practices. Together, pears are among the most cultivated tree fruit crops, with a global yield of 26.5 million tons, providing a rich source of dietary fiber, vitamins, and minerals to temperate regions globally (Reiland and Slavin, 2015; Li et al., 2016; Food and Agriculture Organization of the United Nations, 2024). The sustainability of European pear cultivation has been called into question due to increasing disease pressure, climate change in core growing regions, and the necessity of consistent high yield for economic viability; many of these issues stem from the reliance on cultivars that are over 100 years old (Musacchi, 2024). Consequently, European pear production has declined in the Americas and Europe over the past 20 years (Food and Agriculture Organization of the United Nations, 2024). Asian pear production continues to increase but disease pressure, physiological disorders, and abiotic stresses remain issues in cultivation (Xu et al., 2008; Liu et al., 2013; Zhang and Cui, 2023). Understanding the genetic basis of pear traits is necessary for the development of new cultivars and management practices that can increase the economic sustainability of pear production.

Single linear reference genomes provide the basis for most modern genomics, serving as the foundation for genotyping and resequencing experiments, which can facilitate the connection of genetic variants to phenotypes (Worley et al., 2017). Single-genome references are likely to miss a significant proportion of common variants and loci within a population, however, potentially leading to a failure of sequencing reads to accurately map (Kidd et al., 2007; Miga and Wang, 2020; Eizenga et al., 2025). This lack of whole-genome context contributes to the poor transferability of polygenic scores, as distant and seemingly unrelated loci may be integral to the development of phenotype (Boyle et al., 2017; Mathieson, 2021). This issue leads to considerable problems in the genomic analysis of humans, a species possessing far less genetic diversity than *Pyrus* communis alone, as well as most tree fruit species (Sabety et al., 2024). This contributes in part to the contradiction that pears are a crop of enormous global commercial relevance, yet the genetic basis of most pear traits remains poorly understood.

While many quantitative trait loci have been identified, the lack of stability across genetic backgrounds is a significant limitation to their application in genetic improvement (Iwata et al., 2013; De Mori and Cipriani, 2023). *Pyrus* species also hybridize readily, and many cultivars are the product of crosses between the highly divergent cultivated European and Asian pears (Kumar et al., 2017; Erfani-Moghadam and Zarei, 2018). Assessment of this germplasm is challenging without a reference that includes information from across the genus. The need for genomic representation in wild *Pyrus* species has also been identified, with only a small minority of *Pyrus* species having genomic assemblies (Waite et al., 2024). *Pyrus* species span a wide variety of environments across Afro-Eurasia in cultivated and uncultivated contexts (Figure 1), providing an opportunity to discover genes and variants that contribute to adaptation to diverse climates.

**Figure 1.**
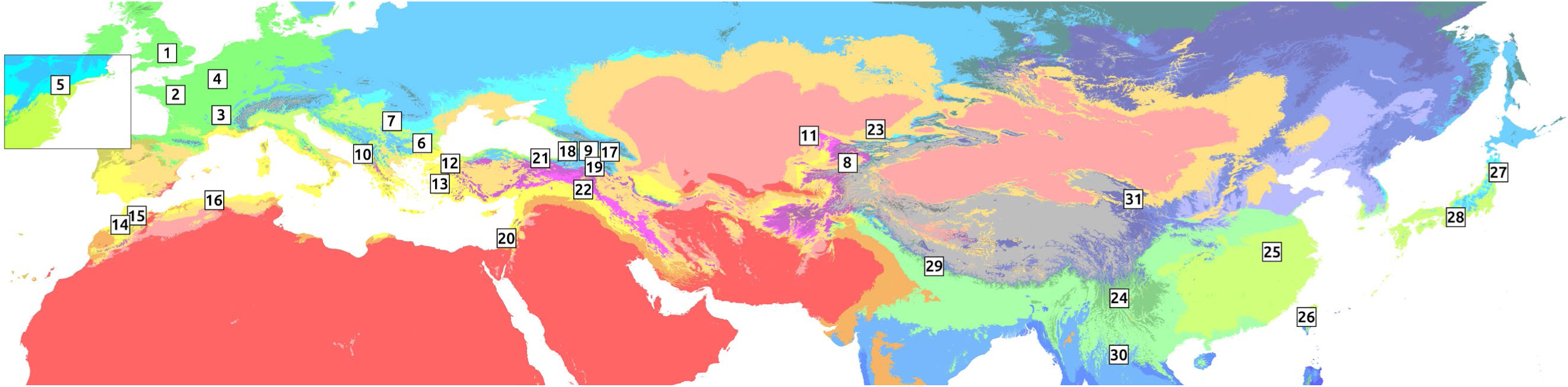
Map showing the origin of sequenced accessions. Insert at left of map depicts the Mid-Atlantic region of the Eastern United States. Background map is the Koppen Climate map for 1991-2020 generated by (Beck et al., 2018). 1– *P. communis* cv. ‘Bartlett’, 2-*P. communis* cv. ‘Comice’, 3– *P. communis* cv. ‘Anjou’, 4– *P. communis* cv. *‘*Abbe Fetel’, 5– *P. communis* cv. ‘Seckel’, 6– *P. communis* cv. ‘Mustafabey’, 7-P. *hybrid* cv. ‘Para de Zahar de Bihor’, 8– *P. korshinskyi*, 9– *P. caucasica*, 10– *P. pyraster*, 11– *P. nivalis*, 12– *P. spinosa*, 13– *P. cordata*\*, 14– *P. gharbiana*\*, 15– *P. mamorensis*\*, 16– *P. cosonii*\*, 17– *P. sachokiana*, 18– *P. salicifolia* (PI657921), 19– *P. salicifolia* (PI312149), 20– *P. syriaca*, 21– *P. elaeagrifolia*\*, 22– *P. glabra*\*, 23– *P. regelii*\*, 24– *P. pseudopashia*\*, 25– *P. calleryana*\*, 26– *P. koehnei*, 27– *P. dimorphophylla*, 28– *P. hondonensis*, 29– *P. pashia*, 30– *P. pyrifolia*, 31– *P. xerophilia*\*. Where precise information on the accession background is unavailable, a representative location for the species was selected – these accessions are marked with an asterisk.

As climate resilience and disease resistance are among the traits most necessary for the sustainability of European pear cultivation (Musacchi, 2024), and these traits are likely to vary widely in response to geographic conditions, these wild germplasm resources are potential sources for genetic variants, which can be of use in genetic improvement efforts, whether by breeding or by gene editing. While attempts have been made to incorporate wild Pyrus germplasm into breeding programs (Claveria et al., 2007, 2012; Dolcet-Sanjuan et al., 2008), the lack of a sufficient genomic reference hinders these efforts and limits the understanding that can be developed from them. For these reasons, the existing genomic references are insufficient for studies aimed at understanding the genetic basis of traits in *Pyrus*. Pangenome graphs solve many of the issues inherent to single reference genomes, while also providing a resource for the discovery of genetic variants of interest (Miga and Wang, 2020; Eizenga et al., 2025). Pangenome graphs improve the accuracy of variant calling, allowing the discovery of structural variants that have previously been uncharacterized (Schreiber et al., 2024; Schloissnig et al., 2025). As advances in sequencing technology enable a greater number of genomic assemblies, it also helps solve the issues posed by the quadratic increase in pairwise comparisons between genomes as new genomes are added to the analysis, which can otherwise hinder interpretability. The development of new tools for the visualization, analysis, and usage of pangenome graphs as references further makes this approach an appealing evolution of traditional genomic analysis (Du et al., 2025). Among tree fruit species, to date, only *Citrus* and *Malus* have received genus-scale pangenome analyses (Huang et al., 2023; Li et al., 2025).

Considering the need for improved genomic references and understanding of adaptive variation in *Pyrus*, alongside the ability to apply pangenome graphs productively, we produced a pangenomic resource for *Pyrus*. To this end, nanopore whole-genome sequencing was applied to 31 Pyrus accessions collected from the USDA National Clonal Germplasm Repository in Corvallis, OR, representing 23 different species. These assemblies were orthogonally validated and then used to generate a pangenome graph using Minigraph.

## Methods

### Sample Collection and Preparation

Fresh leaf tissue from each of 31 accessions was immediately frozen using liquid nitrogen after collection. All samples were cryogenically milled using a Spex Sample Prep Freezer/Mill 6875 (Metuchen, NJ), at a rate of 15cpm for 2 minutes. A two-step protocol was employed for total DNA isolation, first using a modified CTAB DNA isolation protocol, followed by CsCl_2_ density gradient centrifugation for secondary cleanup. Following crude DNA isolation using a modified CTAB protocol, precipitation in equal volume isopropanol, and two washes in 75% ethanol, isopropanol precipitated DNA was resuspended in 5mL 1.56g/mL CsCl_2_ solution at 42°C overnight to ensure complete resuspension. A total of 5µg Ethidium Bromide or 1 µL 10,000X SYBR Gold was added to the sample, loaded into Eppendorf 5PP seal tubes, and brought within 100mg of one another before thermal sealing and centrifugation. The resuspended crude DNA sample was ultracentrifuged for 6 hours at 70,000rpm in a Himac P100VT rotor (Himac, Hitachinaka, Japan). DNA was recovered from the 5PP seal tubes using an 18-gauge needle. The extracted samples were diluted with three volumes of sterile MilliQ (Millipore Sigma, Billerica, MA, USA) water and precipitated with an equal volume of isopropanol. The DNA pellet was washed twice with 75% ethanol, and the samples were resuspended in sterile 1X TE (10mM Tris-HCl, pH 8.0, 1mM EDTA, pH 8.0) overnight at 42°C. Subsequently, the Oxford Nanopore Short Fragment Exclusion Kit (Oxford Nanopore, Oxford, United Kingdom) was used to remove short DNA fragments using the prescribed protocol except for extending the main centrifugation step to 45 minutes from 30 minutes.

### Genomic Sequencing

Isolated DNA samples for each accession were processed for sequencing using the Oxford Nanopore Native Barcoding 24 kit SQK-NBD114.24 (Oxford Nanopore, Oxford, United Kingdom) following the manufacturer’s protocol except for the doubling of enzyme incubation times. 50-100ng of prepared genomic library were loaded into Oxford Nanopore R10.4.1 flow cells and placed into an Oxford Nanopore Promethion P2 Solo device (Oxford Nanopore, Oxford, United Kingdom). Flow cells were washed between sequencing experiments using the Oxford Nanopore Flow Cell Wash Kit (Oxford Nanopore, Oxford, United Kingdom).

### Genomic Assembly

Basecalling of raw .pod5 files was completed using Dorado super accurate basecalling with a quality score cutoff of Q10, followed by demultiplexing and adapter removal using Dorado demux. Seqkit v2.9.0 was used to split .fastq files for each accession into short (3000-62500bp) and long (≥62501bp) read .fastq files (Shen et al., 2024). Short reads were corrected using Dorado correct using default parameters, producing corrected reads in .fasta format. Assembly was performed using Hifiasm v0.25.0, with corrected short reads used as hifi input, and uncorrected long reads being used as ultralong input using the “--ul” parameter (Cheng et al., 2021). The primary contig assembly was used for comparative genomics analysis, while haplotype assemblies were set aside for pangenome construction. RagTag v2.1.0 was used to assign chromosome numbers to contigs and scaffold chromosomes that were not assembled as a full contig (Alonge et al., 2022).

### Orthogonal Validation

Assembly completeness was assessed using BUSCO (version 6.0.0) in genome mode using the eudicots_odb10 reference (Simão et al., 2015). Assembly correctness was assessed using Merqury (version 1.3) (Rhie et al., 2020). Uncorrected and corrected 3000-62500bp reads were assessed using Genomescope (version 2.0) with default parameters to quantify heterozygosity and error rate in both samples (Vurture et al., 2017). Telomere presence was assessed using Telomere Identification Toolkit (Version 0.2.65), with the telomere sequence “AAACCCT” in the 10000bp at the ends of the contigs being used to identify the presence of telomeres (Brown et al., 2025). Telomere sequences inside chromosomes (identified as >50 telomeric repeats in a non-terminal location) were manually split from contigs. Genome Annotation In order to predict genes in primary assemblies, FASTA files were processed using ANNEVO v2.0 for the creation of a gene annotation file (Ye et al., 2025). Annotations were converted into mRNA, CDS, and protein FASTA files using gffread (version 0.12.7) (Pertea and Pertea, 2020). Functional annotation of genes was carried out by two methods: 1) using DeepGO-SE (version 1.0.0) on protein FASTA files using default parameters, and 2) using BLASTP (version 2.16.0) on all predicted protein FASTA files using a database constructed from the *Arabidopsis thaliana* TAIR10 annotation retrieved from Ensembl genomes (Camacho et al., 2009; Kulmanov et al., 2024).

### Comparative Genomics

Colinearity of representative genomes (*Pyrus communis* cv. ‘Bartlett’, *Pyrus mamorensis*, *Pyrus glabra*, *Pyrus regelii*, *Pyrus koehnei*, *Pyrus pyrifolia*, and *Pyrus pseudopashia*) was assessed by D-GENIES (version 1.5.0) (Cabanettes and Klopp, 2018). In order to detect enrichment of specific gene functions in *Pyrus* genomes, annotated proteins from each accession were compared to the combined genes of all accessions using Fisher’s exact test to determine the significance of enrichment. Obsolete and non-plant-related GO terms (e.g., neurogenesis) were manually removed from the final figure.

### Pangenome Graph Construction

The haplotype assemblies from the Hifiasm assembly stage were used for pangenome graph construction. The assembly of the Chinese pear ‘Danxiahong’ was used as a reference/first assembly added to the graph due to its resolution of all telomeric regions (Zhang et al., 2025). Minigraph was used for pangenome graph construction, with default graph assembly parameters (Li et al., 2020). Visualization and graph statistics were generated using the odgi toolkit (Guarracino et al., 2022).

### Genomic Sweep Analysis

*Pyrus* lineages were organized into four groups: all of *Pyrus* subg. *Pashia*, a *P. communis* group, a *P. communis* wild relatives group, and an arid-adapted group. Nucleotide diversity (expressed as pi statistic) and Tajima’s D statistic were calculated for 50kb windows of the genome for each group. Regions of low genomic diversity (D<-1.5) and low variation (pi<10^th^ percentile) were selected as potential selective sweep sites. Genes falling inside these windows were extracted.

## Results and Discussion

### Assembly Performance

A total of 31 highly contiguous genome assemblies were produced, with most chromosomes being assembled in one telomere-to-telomere contig. Within assemblies, 30 to 32 telomeric regions (out of 34) were assembled, with chromosomes 1, 13, and 16 being the most common chromosomes to lack assembled telomeric sequences (Table 1). Summary statistics for all sequenced accessions are provided in Table 1. BUSCO assessed completeness was high; a consistent ∼40% of the assessed conserved orthologs were listed as “duplicated”, a result of the recent whole genome duplication event, which is prominent in the genetic history of Malinae (Velasco et al., 2010; Wu et al., 2013; Xiang et al., 2017). Heterozygosity was high across most tested accessions, with an average of about 1.5% across all accessions (Figure 2a). The assembled genome sizes varied by approximately 10%, from ∼490 Mb to ∼540 Mb across accessions (Figure 2b). No significant differences were detected between the subgenera of *Pyrus*, however the largest and smallest assemblies, as well as the least heterozygous accessions were from *Pyrus* subg. *Pyrus*.

**Figure 2.**
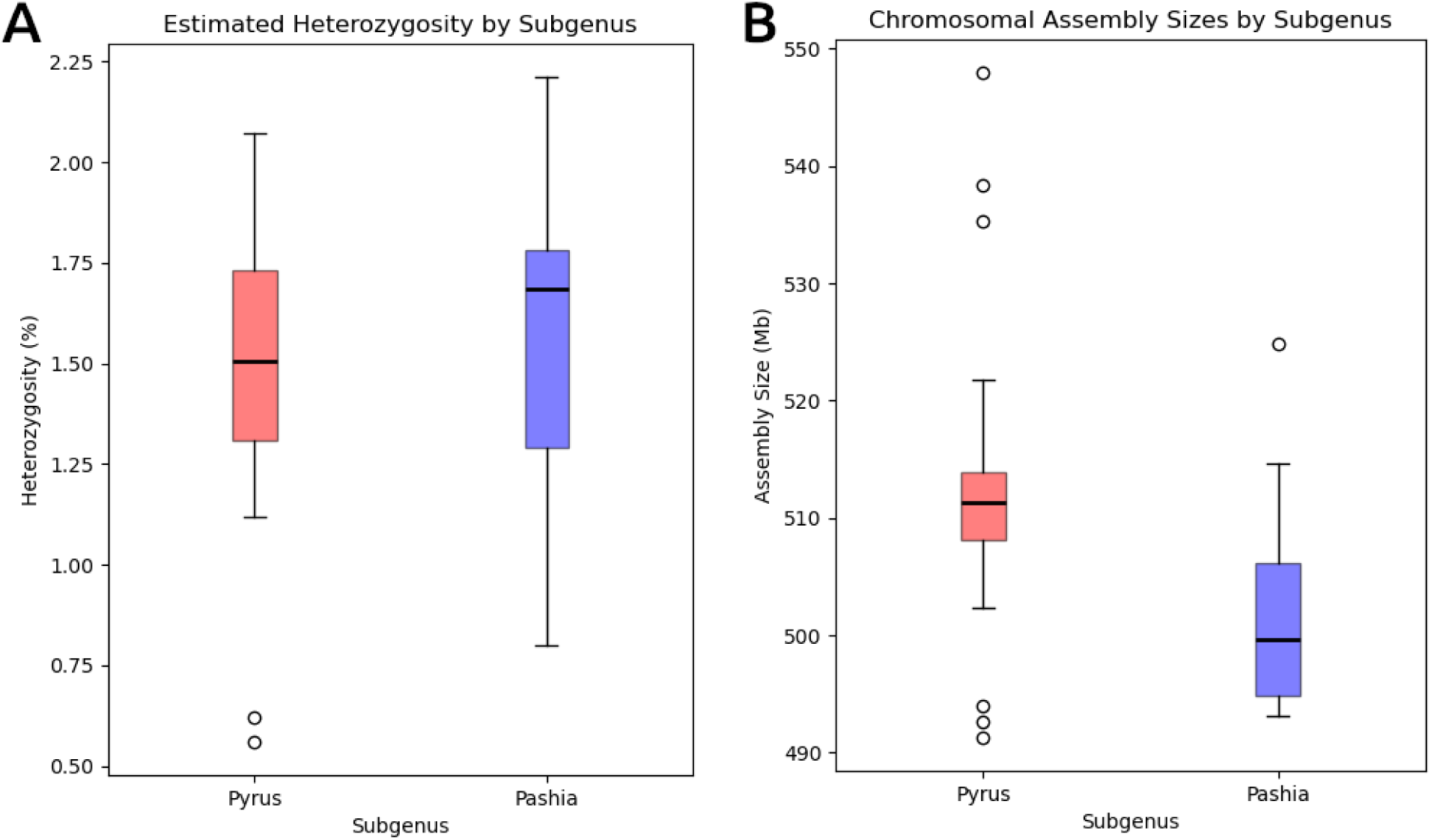
Boxplot graphs displaying. a) estimate heterozygosity and b) total chromosomal assembly sizes for *Pyrus* subg. *Pyrus* and *Pyrus* subg. *Pashia* accessions. No significant differences were found between the groups.

**Table 1.**
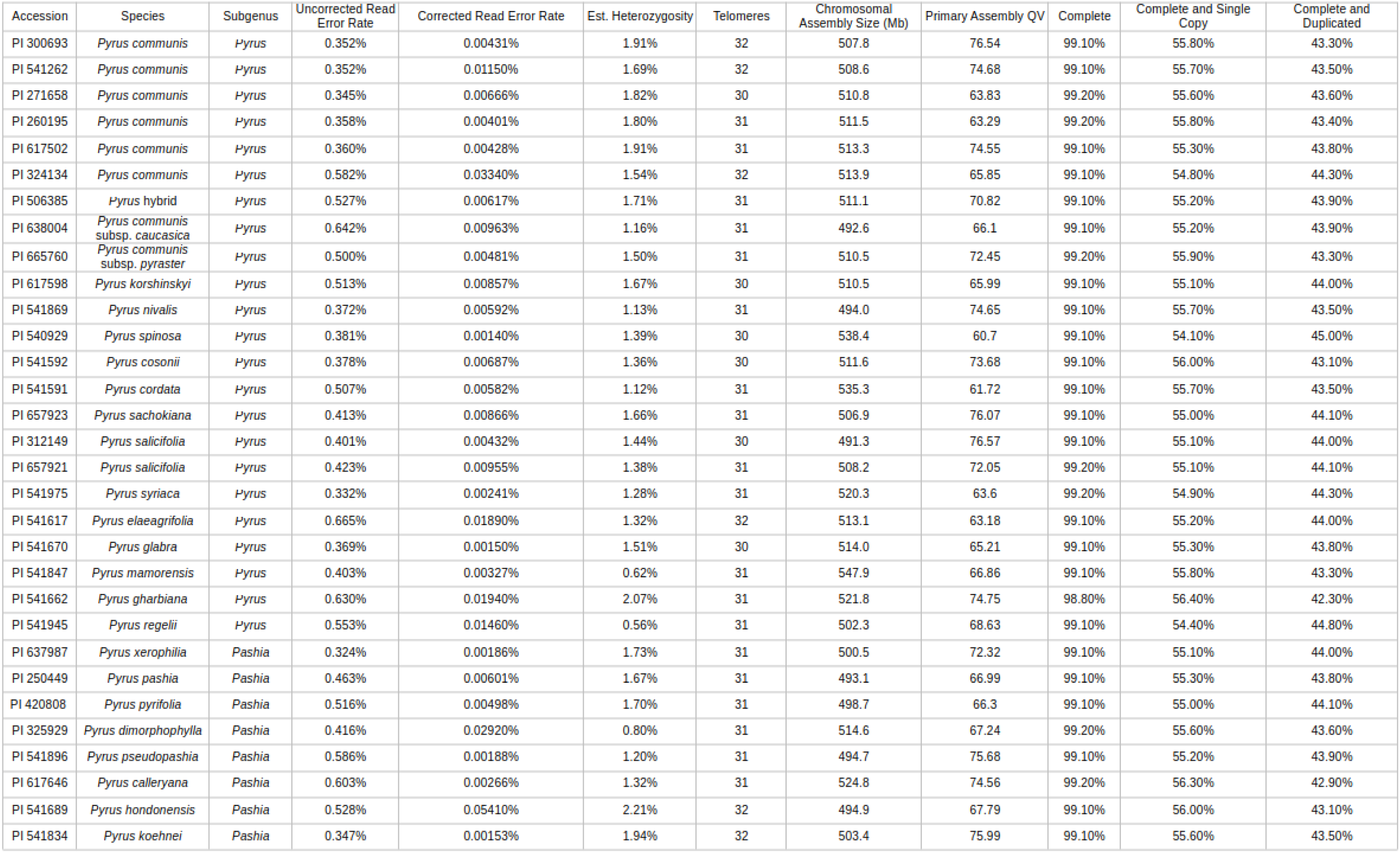
Summary statistics for all accessions sequenced in this study and their primary assemblies.

### Phylogenetic Relationships

While the highly reticulate evolutionary history of *Pyrus* complicates analysis of phylogenetic relationships, inferred phylogeny from protein sequence divergence was largely in agreement with the findings of (Montanari et al., 2020), which grouped *Pyrus* into five general groups: 1) *Pyrus communis*, 2) *Pyrus communis* wild relatives, 3) Middle East arid-adapted species, 4) East Asian pea pears, and 5) East Asian large fruited pears. A *Pyrus communis* grouping was apparent, *Pyrus* subg. *Pyrus* and *Pyrus* subg. *Pashia* were cleanly divided, and an arid-adapted group separate from *Pyrus communis* and its wild relatives was resolved (Figure 3). The *Pyrus communis* subsp. *caucasica* accession was grouped with *Pyrus communis* wild relatives, while *Pyrus korshinskyi* was grouped with *Pyrus communis*, contrary to the groups proposed by (Montanari et al., 2020), although it was noted that *Pyrus korshinkyi* clustered poorly and may not constitute a true species. The placement of *Pyrus regelii* as an outgroup to the rest of the *Pyrus* subg. *Pyrus* groups support prior observations, which suggest that it is an early diverging *Pyrus* lineage that has remained distinct. The *Pyrus regelii* accession has the lowest estimated heterozygosity of all sequenced accessions (Table 1, Figure 2), supported the relative genetic isolation of this species.

**Figure 3.**
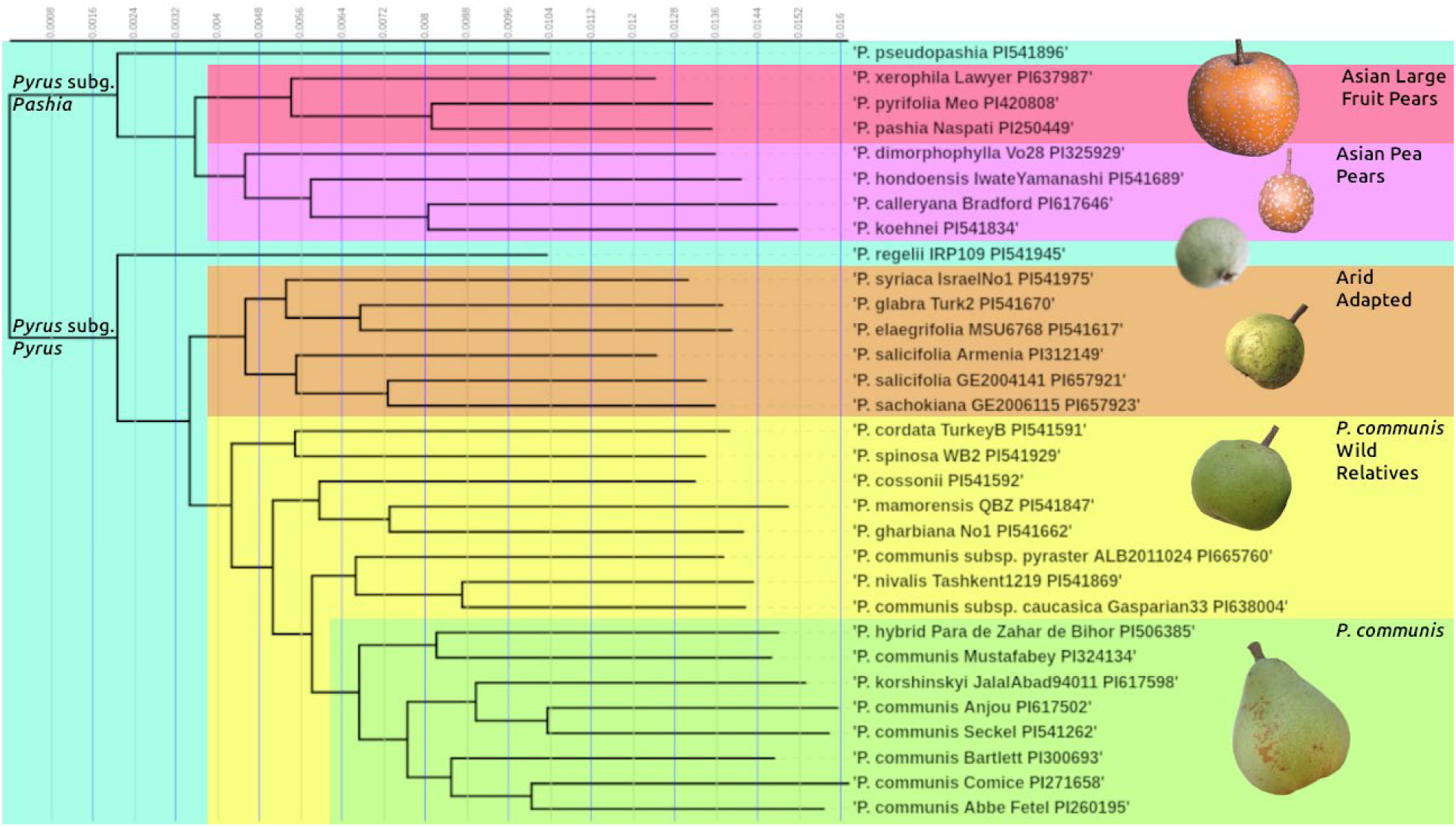
Species tree inferred by protein similarity. Colored blocks are placed around the 5 previously identified groups (labelled at right of figure) within *Pyrus*. *P. regelii* and *P. pseudopashia* appear to be basal lineages of *Pyrus* subg. *Pyrus* and *Pyrus* subg. *Pashia*, respectively, and are left ungrouped (background in teal).

### Pairwise Comparative Genomics

To explore genomic collinearity between samples, representative samples from each group were taken, 7 in total. This was done to avoid complications arising from the very high number of total pairwise comparisons between 31 genomes (465 comparisons). Collinearity was strong between the representative genomes, with only a few major, megabase-scale alterations to genome structure apparent, such as a large inversion on Chromosome 13 of *P. mamorensis*, Chromosome 6 of *P. communis* cv. ‘Bartlett’, and Chromosome 14 of *P. pseudopashia* (Figure 4). Inversions can be important events in adaptive evolution, as they can reduce recombination of alleles in the inverted regions, and maintain adaptive traits in populations even in the face of the extensive gene flow that is typical of many plant lineages (Farré et al., 2013; Oneal et al., 2014; Twyford and Friedman, 2015; Wellenreuther and Bernatchez, 2018). Within these large, detected inversions, there were numerous genes (107 in *P. mamorensis*, 30 in *P. communis* ‘Bartlett’, and 48 in *P. pseudopashia*) that have potential roles in environmental adaptation. The inversion in *P. mamorensis*, for example, contains what appear to be arrays of tandem-duplicated predicted MLP-like genes (13 predicted genes in a ∼1.1Mb inversion), genes which are known to be responsible for abscisic acid and ethylene regulation, conferring drought tolerance in rice and *Arabidopsis* (Ruperti et al., 2002; Wang et al., 2016; Zhou et al., 2022). These genes are also implicated in both positive and negative regulation of pathogen defense in fruit crops (He et al., 2020; Sun et al., 2023; Su et al., 2024), which in context may relate to climate-dependent pathogen risks. Other putative tandem duplications within these inversions include homologs of TOPLESS-related 2, cytochrome P450 genes, and cupin superfamily genes (Table 2), many of which share stress-response related functions which may drive environmental adaptation (Swaminathan et al., 2009; Höfer et al., 2014; Wang et al., 2014; Griebel et al., 2023). These regions may be worth considering in future genotyping of a wider array of *Pyrus* germplasm.

**Figure 4.**
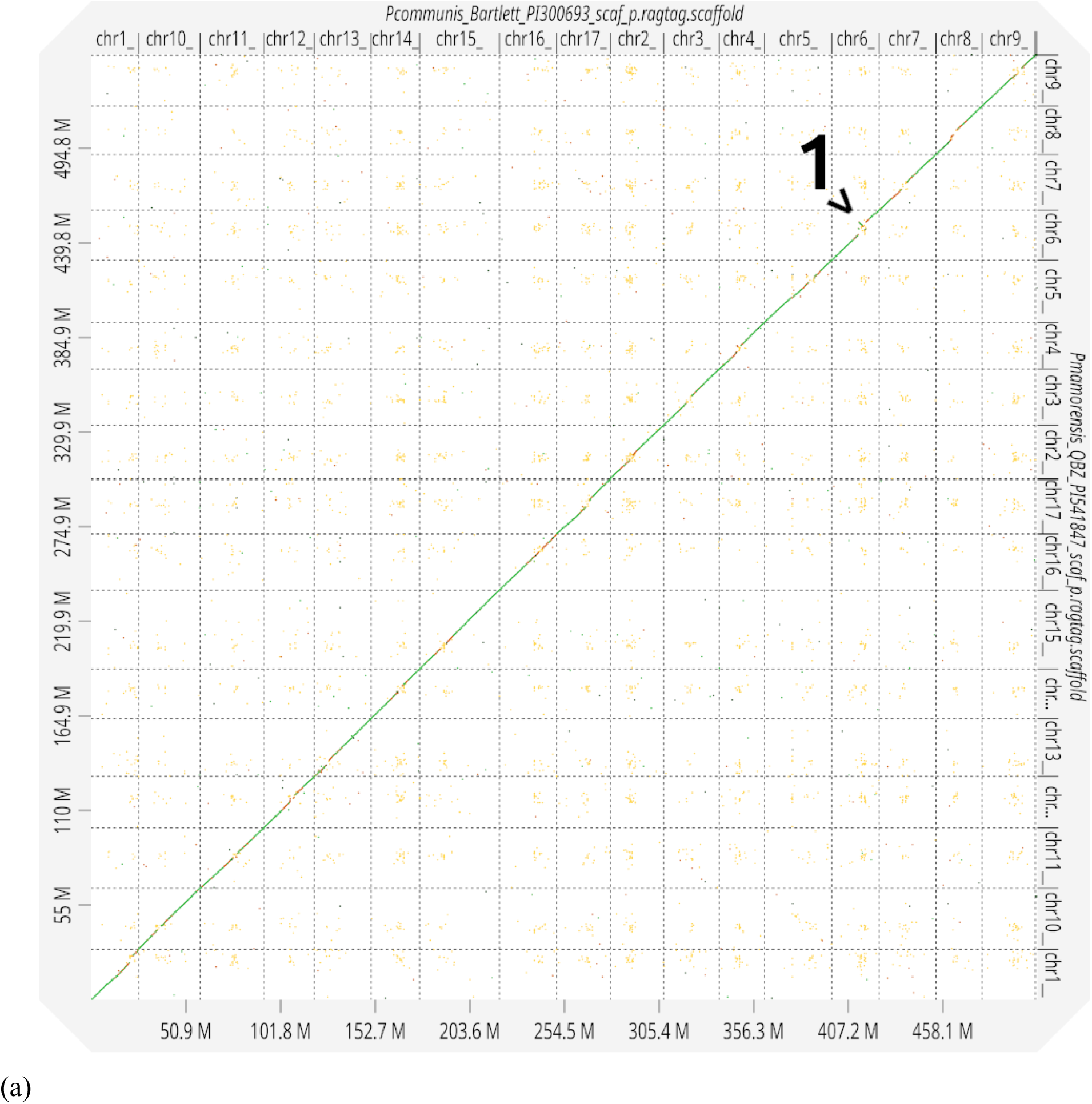
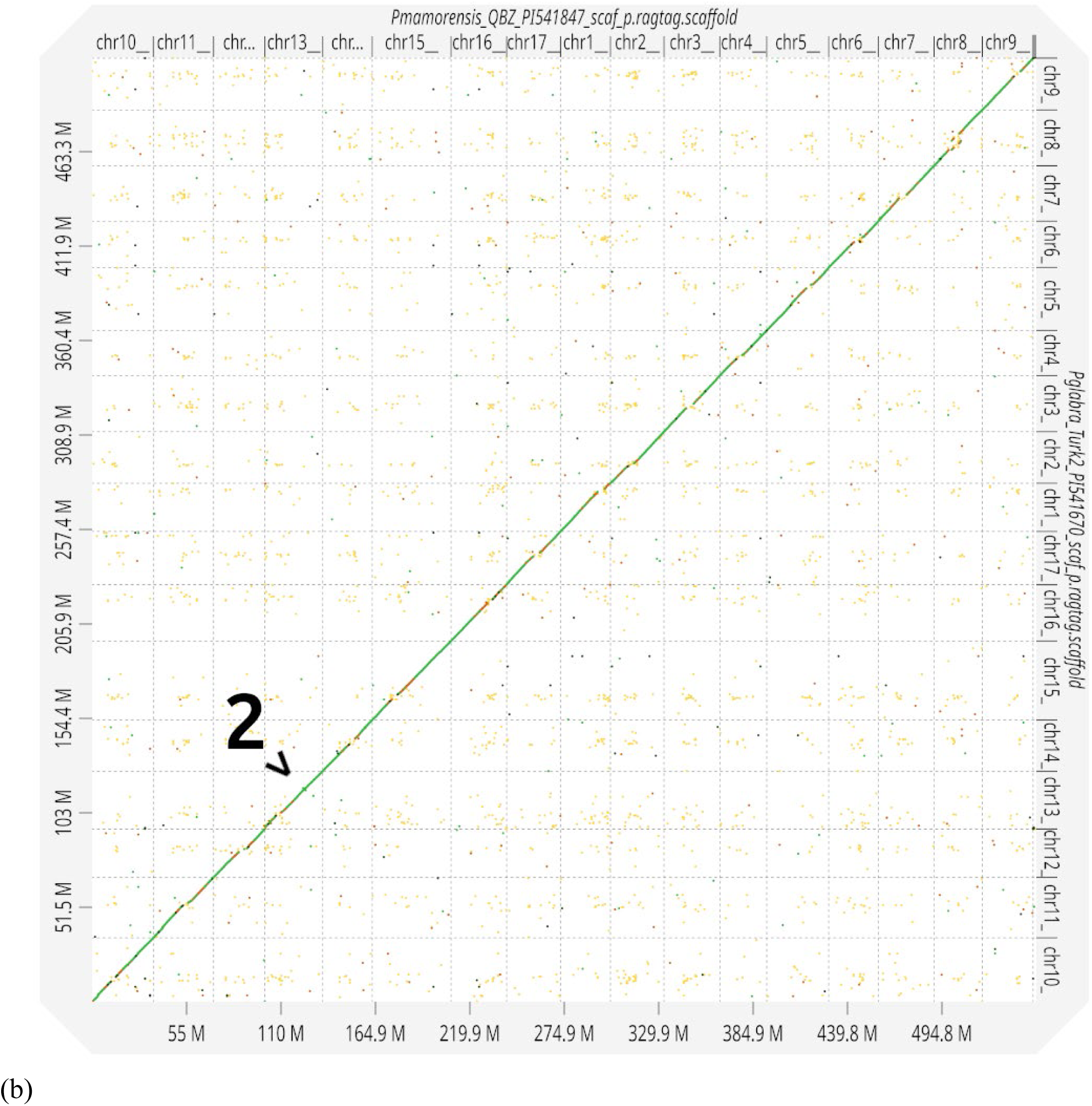
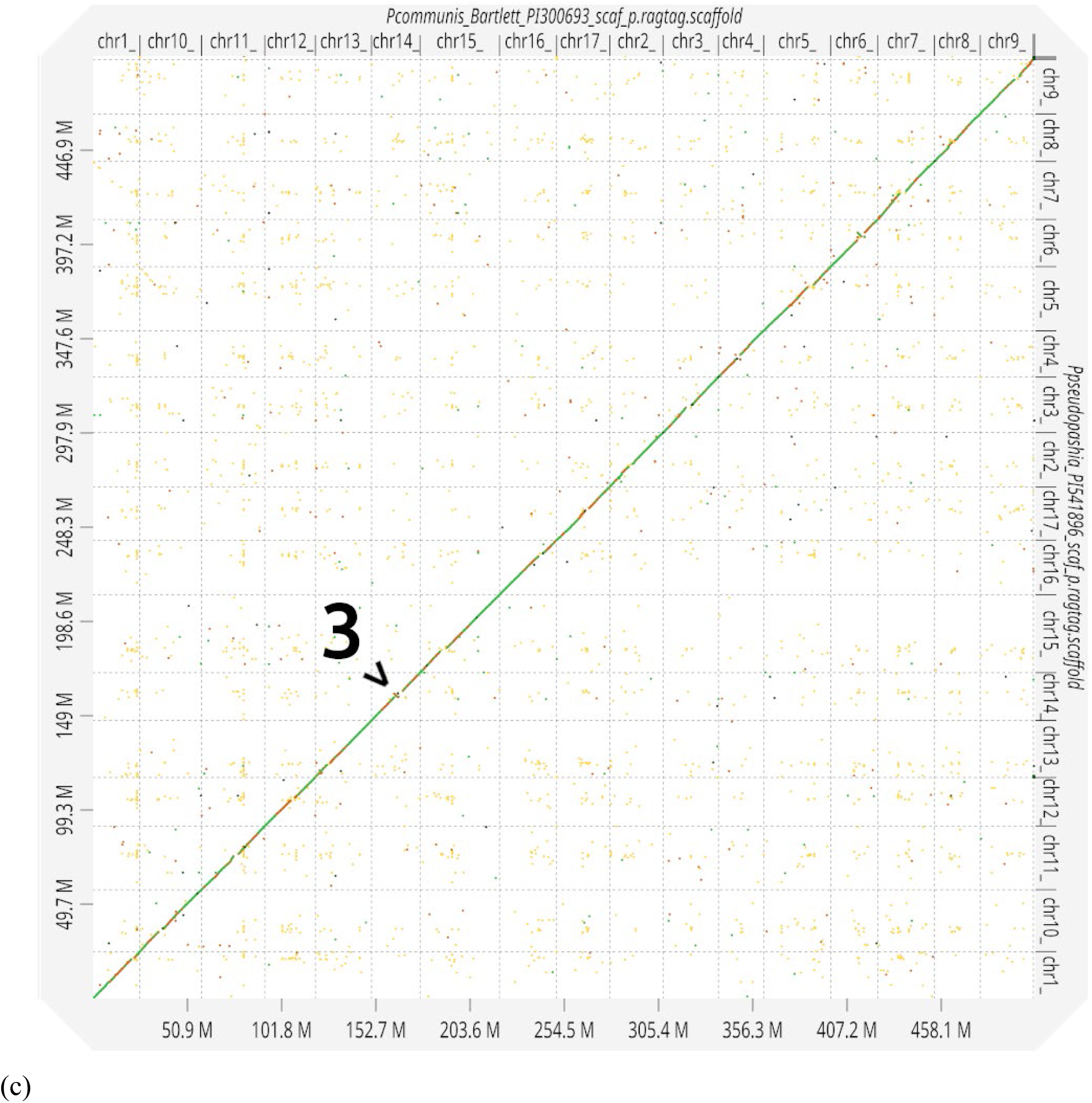
Detection of large inversions by examination of collinearity. Collinearity between (a) *P. communis* and *P. mamorensis*, (b) *P. mamorensis* and *P. glabra*, and (c) *P. communis* and *P. pseudopashia* displayed. Major inversions marked for 1) *P. communis*, 2) *P. mamorensis*, and 3) *P. pseudopashia*.

**Table 2.** Putative tandem repeats within identified large inversions in selected *Pyrus* species. Neighboring genes with different *Arabidopsis* gene homologs with the same annotation were included.

| **Inversion** | **Genes** | **Arabidopsis Homolog** | **Annotation** |
| --- | --- | --- | --- |
| *P. mamorensis*<br>Chromosome 13 | g391.t1, g392.t1 | AT3G16830.1 | TOPLESS-related 2 |
| *P. mamorensis*<br>Chromosome 13 | g395.t1, g396.t1,<br>g397.t1, g398.t1,<br>g399.t1, g400.t1,<br>g401.t1, g402.t1 | AT1G24020.1 | MLP-like protein 423 |
| *P. mamorensis*<br>Chromosome 13 | g1376.t1, g1377.t1,<br>g1378.t1 | AT5G27870.1 | Plant invertase/pectin<br>methylesterase inhibitor superfamily |
| *P. mamorensis*<br>Chromosome 13 | g1390.t1, g1391.t1,<br>g1392.t1, g1394.t1 | AT1G24020.1 | MLP-like protein 423 |
| *P. mamorensis*<br>Chromosome 13 | g1409.t1, g1410.t1 | AT4G02080.1 | secretion-associated RAS super<br>family 2 |
| *P. communis*<br>Chromosome 6 | g639.t1, g640.t1,<br>g641.t1 | AT2G45180.1,<br>AT1G12100.1 | Bifunctional inhibitor/lipid-transfer<br>protein/seed storage 2S albumin<br>superfamily protein |
| *P. pseudopashia*<br>Chromosome 14 | g584.t1, g585.t1 | AT3G05950.1 | RmlC-like cupins superfamily protein |
| *P. pseudopashia*<br>Chromosome 14 | g586.t1, g587.t1 | AT3G52970.1 | cytochrome P450, family 76,<br>subfamily G, polypeptide 1 |

The comparison of predicted proteomes revealed a large core genome. Accessions had between 33,784 and 34,842 orthogroups present individually, while 21,375 orthogroups were determined to be a part of the core genome (Figure 5). While genes belonging to the private genome (only observed in one accession) are very rare, about half of the detected orthogroups fell into the accessory (found in 2-6 genomes) and dispensable (7-29 genomes) groups, showing a wide diversity in the genes present in *Pyrus* species (Figure 6). However, only a small handful of functional categories show significant enrichment or depletion while comparing the subgroups of *Pyrus* (Table 3). Reproduction-related genes showed enrichment across the Asian Pea Pears, while Asian Large Fruit Pears show an elevation in annotations related to polysaccharide metabolism, fatty acid synthesis, and urea transport. Differences between the *Pyrus* subg. *Pyrus* groups were limited as well, with the *P. communis* group showing enrichment in magnesium and redox-related processes, *P. communis* wild relatives showing alterations in amino acid functions, and the Arid-Adapted group showing a notable increase in membrane raft-related genes. It is likely that, owing to the relatively recent divergence of *Pyrus*, functional diversity is achieved through more subtle allelic differences and variation in gene regulation.

**Figure 5.**
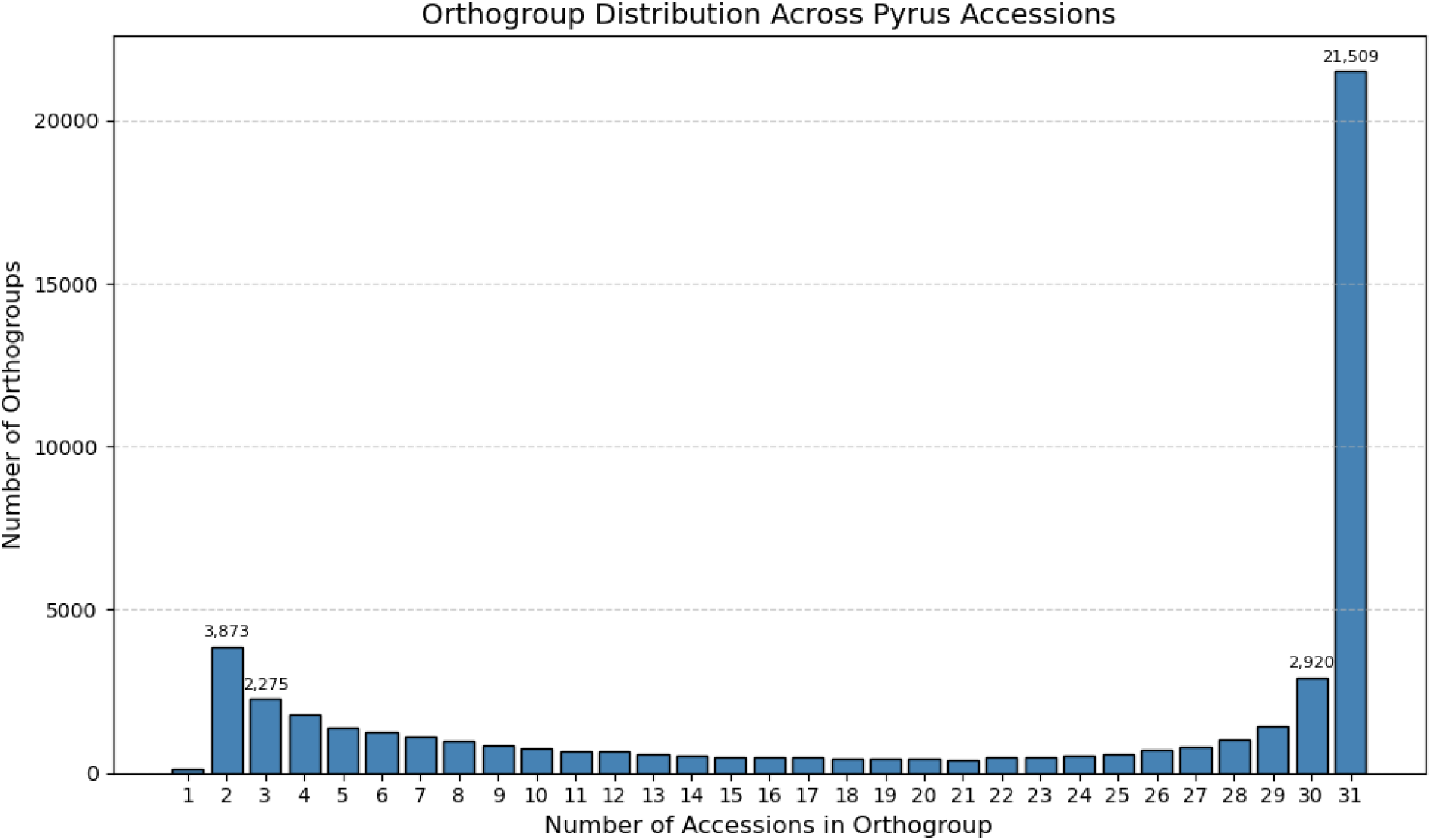
Orthogroup frequency across the 31 *Pyrus* accessions sequenced for this study. Bars indicate the number of orthogroups observed in *k* accessions (x-axis). For example, 2,920 orthogroups are found in 30 of 31 accessions sequenced.

**Figure 6.**
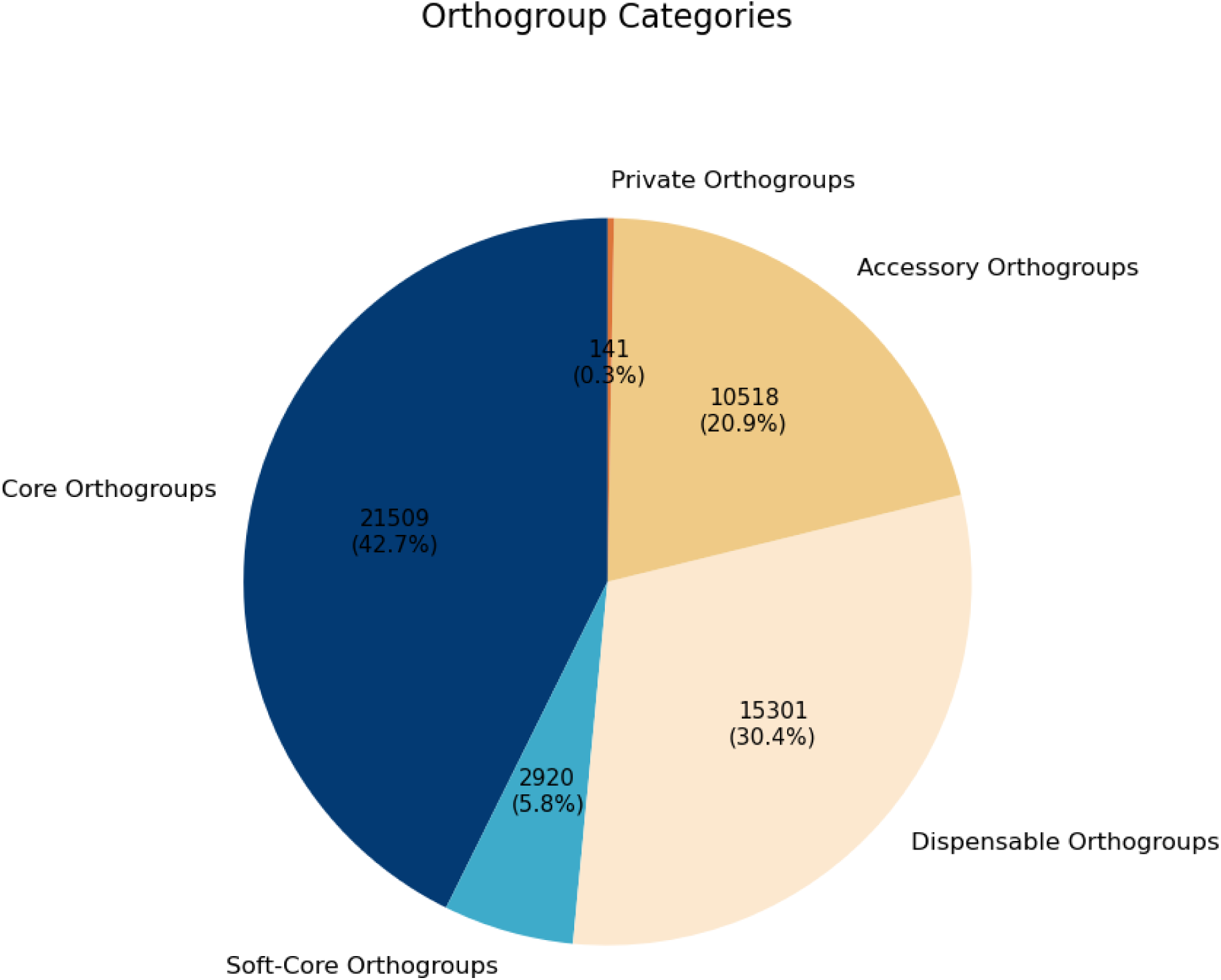
Proportion of detected orthogroups belonging to core (all accessions), soft-core (30 accessions), dispensable (6-29 accessions), accessory (2-5 accessions), and private (1 accession) genomes.

**Table 3.**
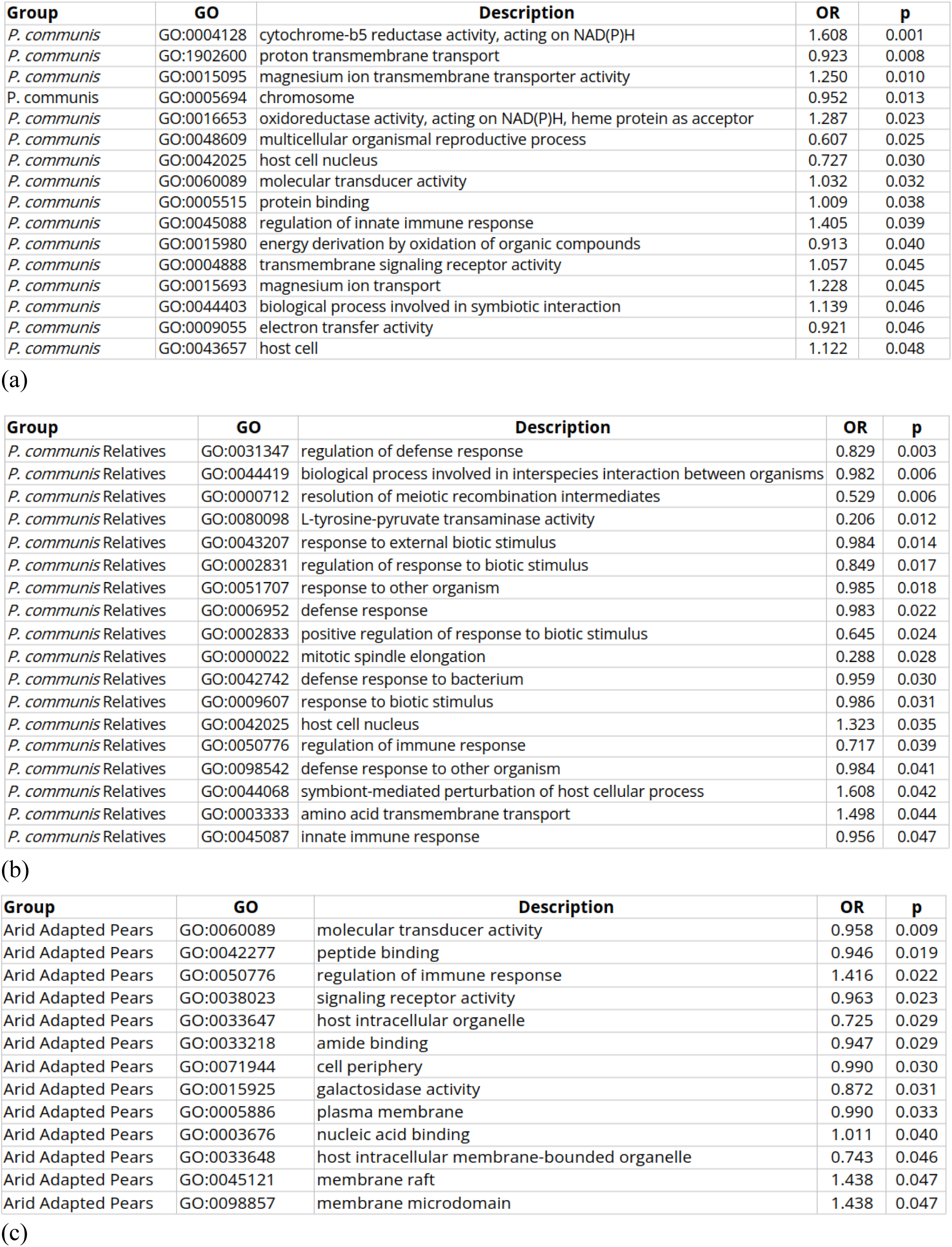
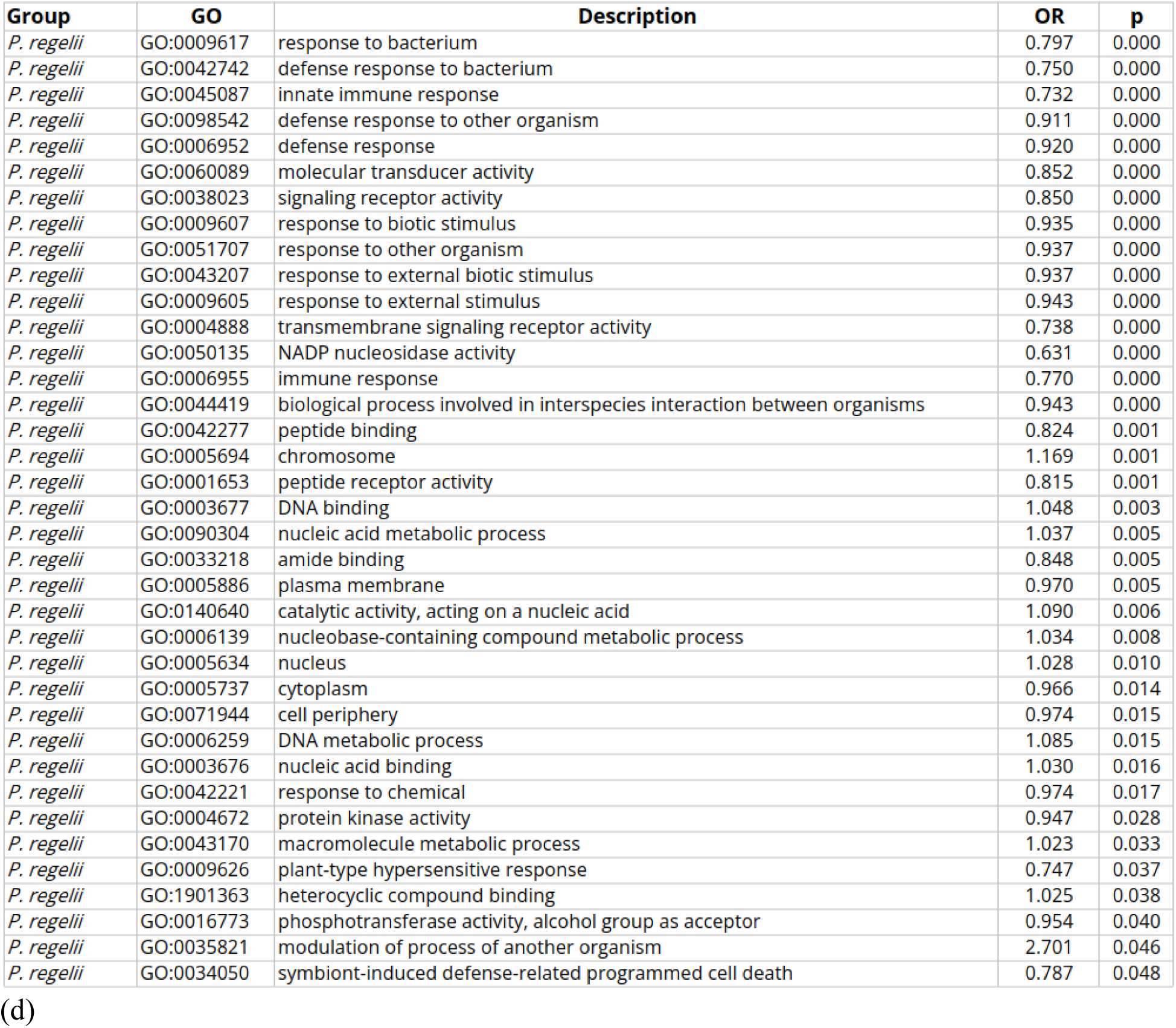
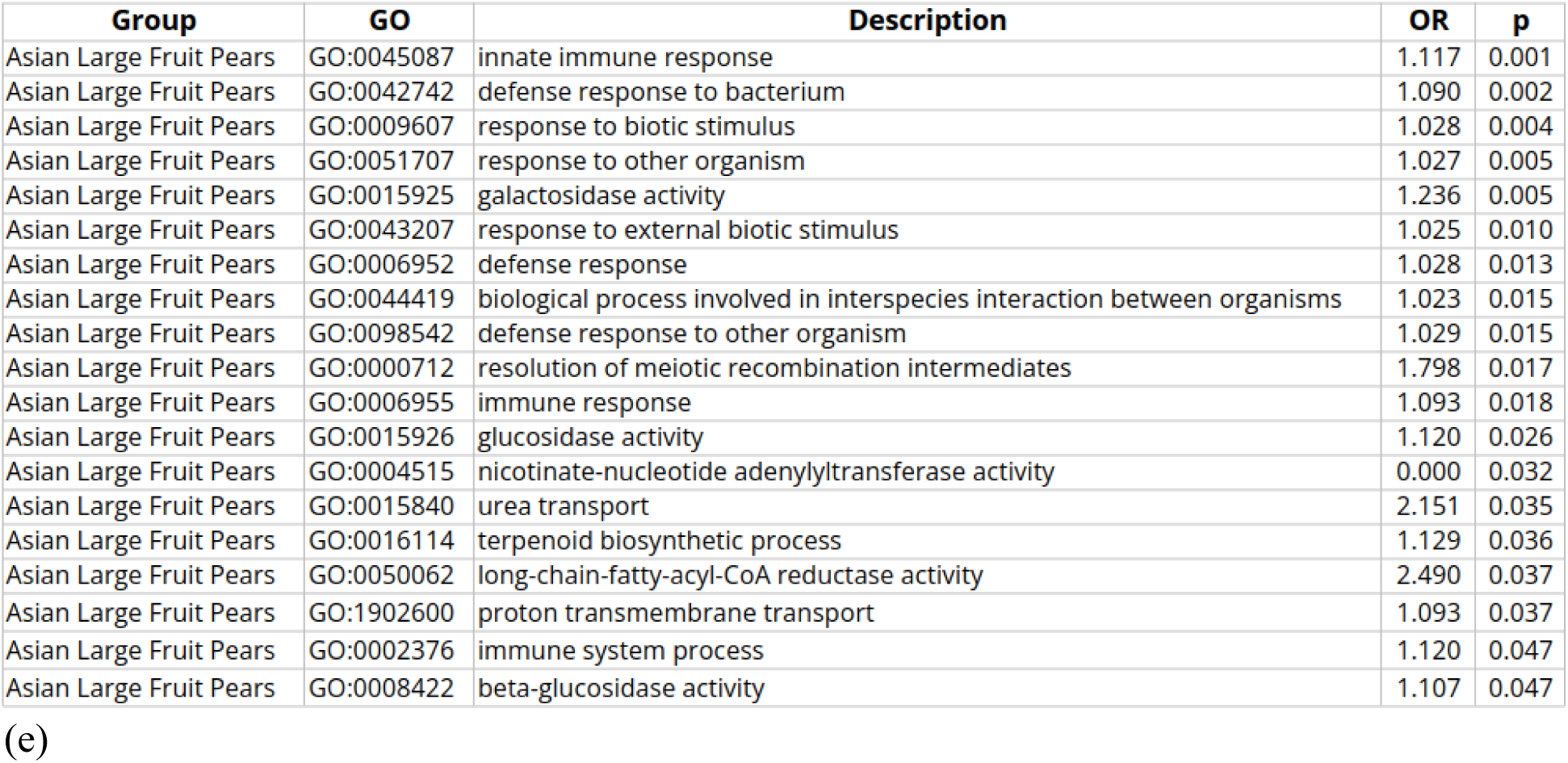
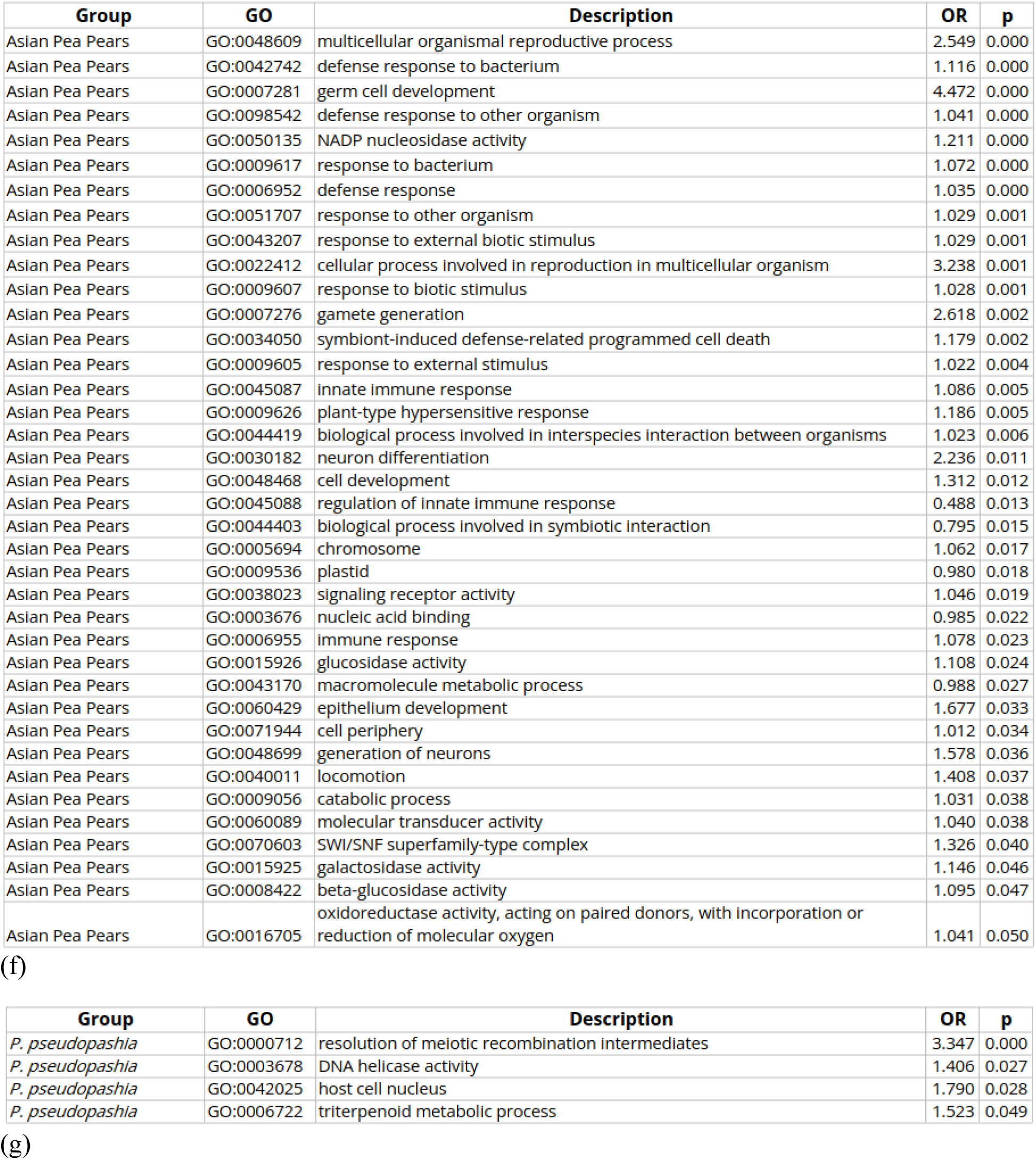
Enriched GO terms in orthogroups unique to different *Pyrus* subgroups. ((a) *Pyrus communis*, (b) *Pyrus communis* Relatives, (c) *Arid Adapted* Pears, (d) *Pyrus regelii*, (e) Large Fruit Asian Pears, (f) Asian Pea Pears, (g) *Pyrus pseudopashia*) at p<0.05.

### Pangenome Graph

To explore the genetic diversity of *Pyrus* in finer detail, a pangenome graph was generated from the selected accessions; the resulting graph contains 2,978,098,326 base pairs, 274,887,871 nodes, and 380,638,055 edges. The sequence contained within the graph is approximately 6x larger than the average haploid genome size of *Pyrus* individuals. The high degree of complexity of the graph is reflective of the high degree of genetic diversity within *Pyrus*. While subdivision of the graph may be appealing (especially along the obvious division between *Pyrus* subg. *Pyrus* and *Pyrus* subg. *Pashia*) to make the graph more succinct and computationally efficient, the extensive gene flow even across this division would weaken the ability of the graph to capture all variation in both subgenera. The graph contains ∼25 million variants, which can be used for genotyping of *Pyrus* samples (Figure 7), generated from across the genus, enabling calling of all variants, including large structural variants, even using short read data (Ebler et al., 2022). A core pangenome of 210.2 Mb could be identified, with most of the sequence of each accession being shared by a smaller group of accessions (Table 4). While previous analysis of NCGR germplasm cast doubts on the taxonomic classification of the *Pyrus pseudopashia* accession PI541896 (Montanari et al., 2020), the high degree of private sequence relative to other *Pyrus* subg. *Pashia* accessions, as well as the positioning as an outgroup to the rest of *Pyrus* subg. *Pashia* by protein sequence identity suggests this accession may be properly classified, representative of this early divergent species within *Pyrus* subg. *Pashia* (Zheng et al., 2014a; Jin et al., 2024b).

**Figure 7.**
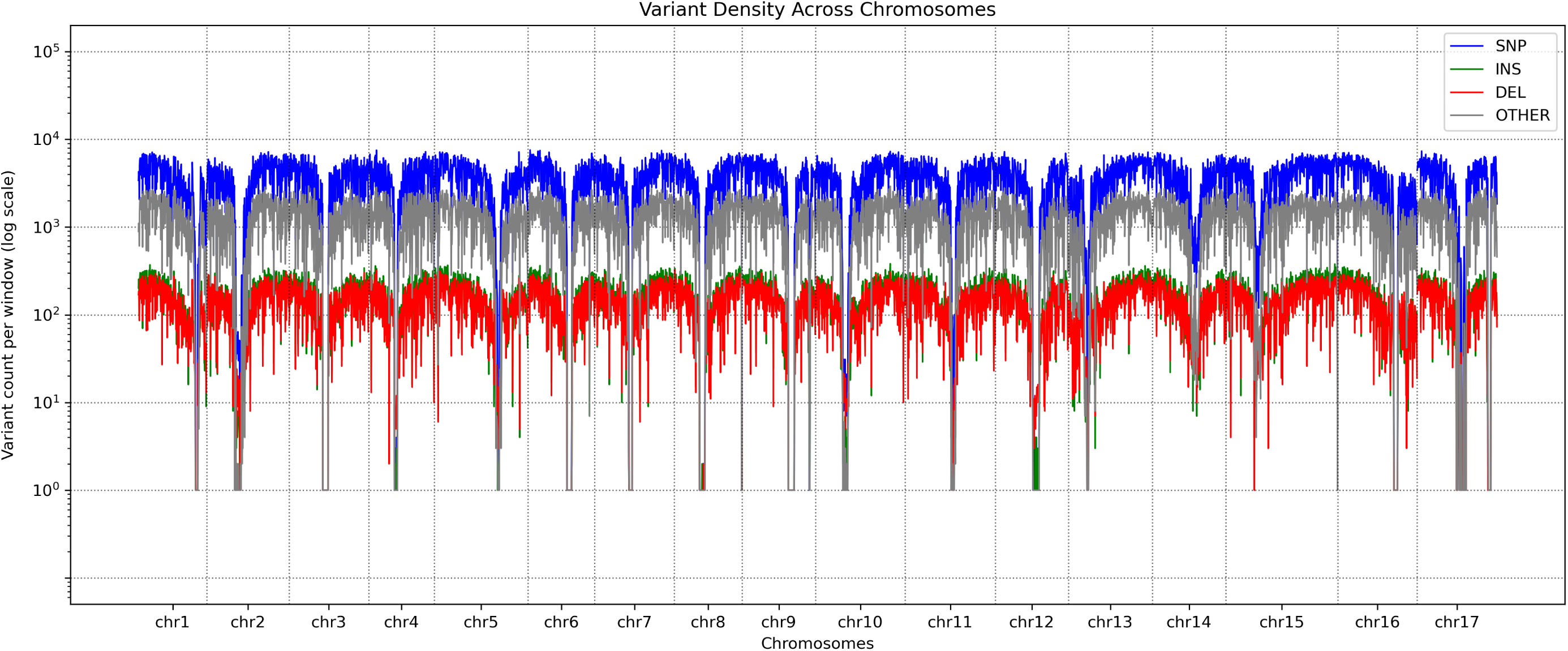
Distribution of genetic variants in *Pyrus* established by Minigraph-Cactus pangenome graph construction of 64 haplotypes. Regions of low detected variation correspond to centromeric regions.

**Table 4.**
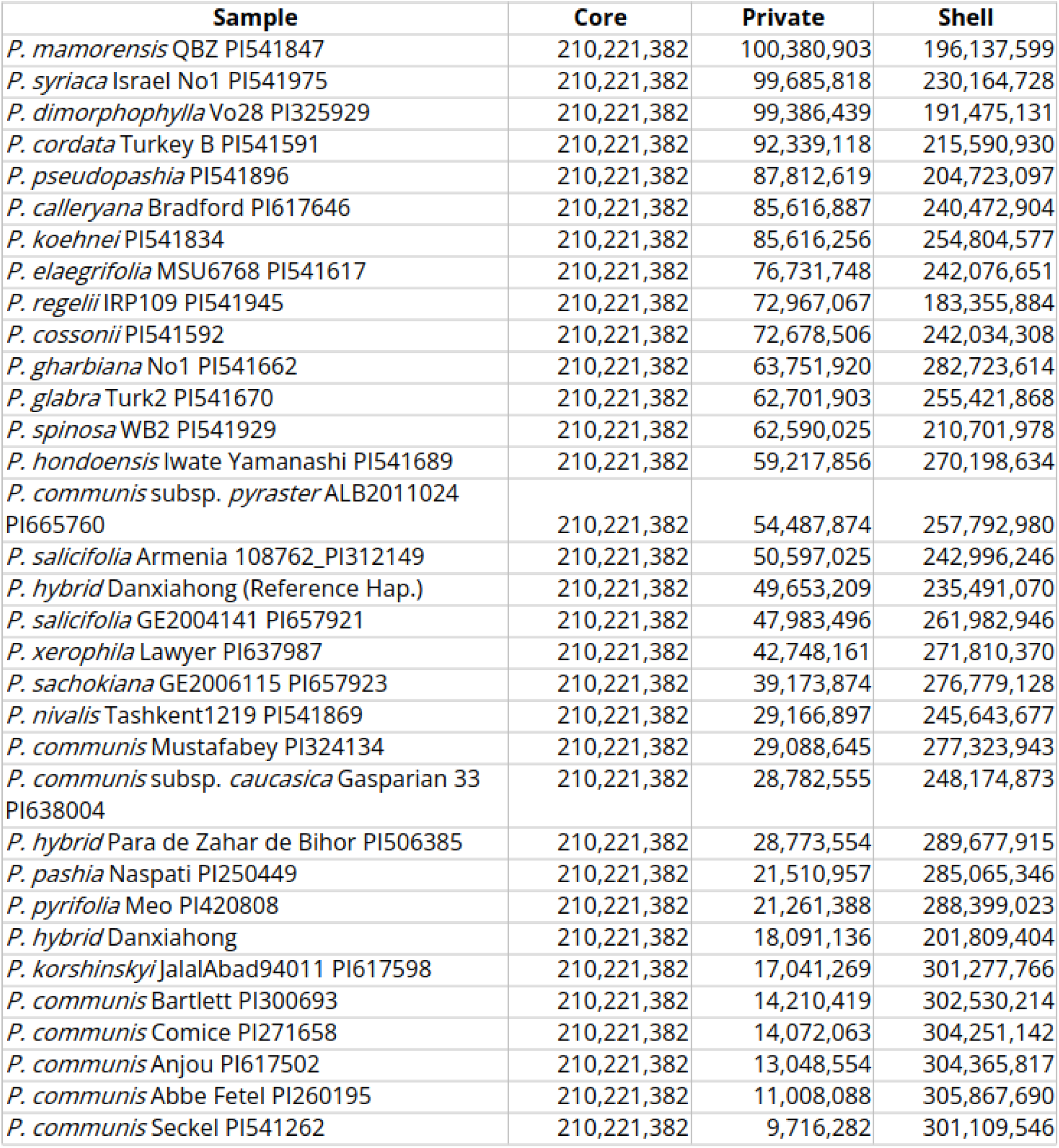
Pangenome size divided into core (shared by all accessions), private (unique to one accession), and shell (shared by some but not all accessions) pangenome.

### Pangenome-Aided Detection of Climate-Related Genes

The group of arid-adapted *Pyrus* species appears as natural candidates in exploring variants that may be responsible for adaptation to abiotic stress resilience. The estimated divergence of *Pyrus* subg. *Pyrus* is relatively recent, in an environmental context of much of Asia becoming more arid (Guo et al., 2004; Caves et al., 2014; Korotkova et al., 2018; Ao et al., 2021), perhaps leaving signs of harsh selective events in the genomes of these species. Potential sites with selective sweeps were identified for each group (Table 5). In the Arid Adapted Group, for example, 12 putative selective sweep windows were identified, with 33 genes falling within these windows, the most of any of the groups. Genes within these regions include ones with known function in abiotic stress resistance (Table 6), for example, an ANR1 homolog, which is responsive to abscisic acid and abiotic stress more broadly (Lin et al., 2020), and flavin-binding monooxygenases which have been implicated in drought and salt stress tolerance (Gaba et al., 2023; Wang et al., 2023; Zhao et al., 2024). While the ANR1 homolog locus shows signs of a selective sweep considering SNP sites, there are considerable structural variations in this region (Figure 8), and this locus appears to be greatly expanded in *Pyrus* subg. *Pyrus* relative to *Pyrus* subg. *Pashia* (Figure 9). Stress response related genes were found to be regions enriched in structural variants in rice, and structural variants, particularly resulting from the movement of transposable elements, were often found even in regions with sweep signals in rice, maize, and poplar (Yang et al., 2013; Wu et al., 2018; Fuentes et al., 2019; Zhao et al., 2022). The fact that many of the genes which fall within putative sweep sites appear to be transcription factors (Table 6), may help explain the recent and rapid divergence of *Pyrus* to varied climates across Afro-Eurasia, especially arid and semi-arid regions. The ability for alterations on regulatory sequences to have profound impacts on phenotype and evolutionary adaptation is well understood, and this principle applies to both cis– and trans-acting regulatory elements (King and Wilson, 1975; Wittkopp et al., 2004; Carroll, 2005; Signor and Nuzhdin, 2018). Transcription factors in particular can regulate the expression of many genes, with alterations potentially having an outsized impact, especially in adaptation to new environments (Saxer et al., 2014). In the *Malus* pangenome, wild *Malus* germplasm possessed a SNP in the promotor region of the MYB5 transcription factor gene, which could be identified as a contributor to cold stress tolerance (Li et al., 2025). While this preliminary analysis suggests an important role for transcription factors in recent *Pyrus* adaptations to arid environments, genotyping a wider array of *Pyrus* individuals, as enabled by this pangenome graph, may allow for higher confidence identification of a wider number of sites that have experienced recent adaptive selection. Pantranscriptomic analysis may be particularly warranted in further exploration of wild *Pyrus* species in their adaptation to arid environments, due to the prominent role of transcription factors discovered in this analysis.

**Figure 8.**
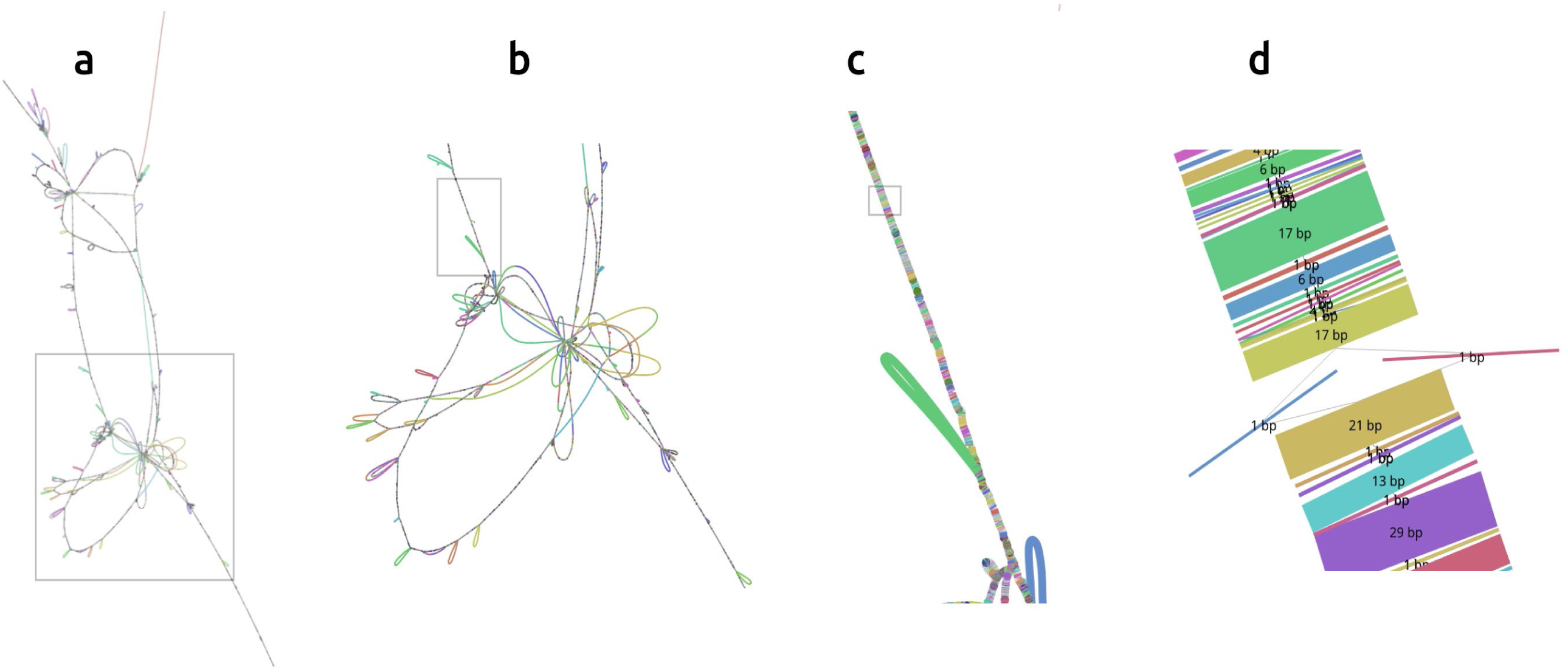
ANR1 locus and vicinity viewed at multiple spatial scales. (a) At large (>50kb) spatial scales only the largest structural variants and regions of high variability are clearly visible. (b) Visualized at a slightly smaller scale, one portion of the ANR1 locus shows many insertions (typically represented as loop-like paths adjacent to the main path) and divergent paths. (c) At a smaller spatial scale (<10kb), smaller variants start to become visible; an 8.6 kb insertion unique to one haplotype of *P. calleryana* (large green node, bottom center) is apparent. (d) A 1bp variant (T->A) found only in one haplotype of *P. communis* ‘Seckel’ (red node), displaying the single base pair resolution of the pangenome graph. Visualization performed using Bandage-NG. Grey boxes indicate the region zoomed in on the panel to the right.

**Figure 9.**
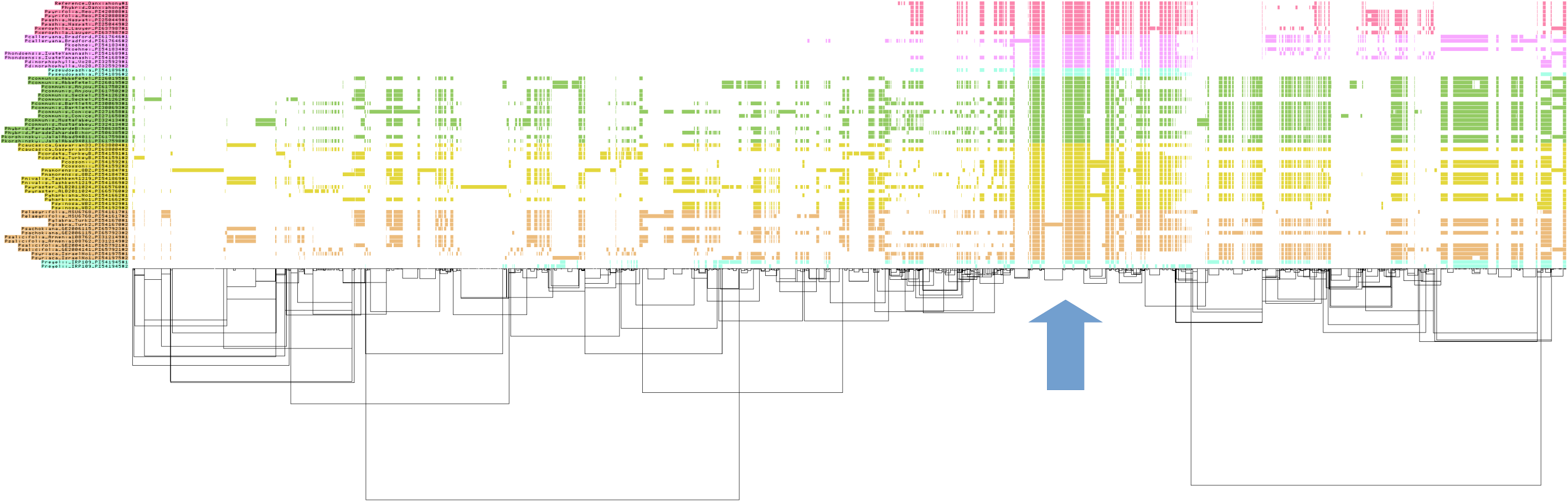
Diagram of presence-absence variants at the predicted ANR1 locus (gene length ± 10kb) with a bin size of 250bp. *Pyrus* subgroups share a color as in Figure 4. Haplotypes of each accession are indicated by #1 and #2 for haplotypes 1 and 2, respectively. Accessions are grouped by color as in Figure 4. Bars in a given row indicate the presence of sequence. A core region shared by most accessions is apparent (denoted by a blue arrow below), with a larger region being shared by most *Pyrus* subg. *Pyrus* accessions.

**Table 5.** Windows showing signs of selective sweeps within *Pyrus* subgroups. Regions satisfy all of the following: >300 SNPs in the window, Tajima’s D statistic <= –1.5, and pi statistic in the 10^th^ percentile or lower.

| Group | Chromosome | Window Start | Window End | PI | Tajima's D Statistic | Number of Variants | Number of SNPs |
| --- | --- | --- | --- | --- | --- | --- | --- |
| Arid | 3 | 6750000 | 6800000 | 0.00279483 | -1.6704 | 569 | 334 |
| Arid | 3 | 16300000 | 16350000 | 0.00243388 | -1.52771 | 505 | 322 |
| Arid | 3 | 18200000 | 18250000 | 0.00268537 | -1.76218 | 607 | 372 |
| Arid | 4 | 19200000 | 19250000 | 0.002402 | -1.7546 | 546 | 343 |
| Arid | 5 | 28950000 | 29000000 | 0.002754 | -1.61918 | 596 | 357 |
| Arid | 6 | 17400000 | 17450000 | 0.00257353 | -1.75532 | 568 | 317 |
| Arid | 7 | 50000 | 100000 | 0.00225683 | -1.8318 | 511 | 313 |
| Arid | 7 | 2850000 | 2900000 | 0.002842 | -1.55608 | 566 | 310 |
| Arid | 7 | 19250000 | 19300000 | 0.00279753 | -1.94221 | 616 | 318 |
| Arid | 13 | 19550000 | 19600000 | 0.00205719 | -1.90189 | 485 | 314 |
| Arid | 16 | 17100000 | 17150000 | 0.00276906 | -1.54316 | 583 | 354 |
| Arid | 16 | 19150000 | 19200000 | 0.0024857 | -1.56813 | 517 | 332 |
| Communis | 10 | 16550000 | 16600000 | 0.00103344 | -1.87492 | 425 | 328 |
| Communis | 11 | 0 | 50000 | 0.00256976 | -1.92613 | 654 | 421 |
| Communis | 16 | 2150000 | 2200000 | 0.002514 | -1.8358 | 624 | 378 |
| Communis | 16 | 4950000 | 5000000 | 0.002581 | -1.76387 | 649 | 307 |
| Communis | 17 | 12000000 | 12050000 | 0.00131711 | -1.96037 | 551 | 364 |
| Communis Relatives | 12 | 0 | 50000 | 0.00270515 | -1.54202 | 533 | 367 |
| Pashia | 1 | 24350000 | 24400000 | 0.00227108 | -1.62614 | 583 | 380 |
| Pashia | 2 | 23200000 | 23250000 | 0.00246551 | -1.60484 | 596 | 316 |
| Pashia | 6 | 11700000 | 11750000 | 0.00216927 | -1.6649 | 554 | 332 |
| Pashia | 10 | 0 | 50000 | 0.0022235 | -1.57709 | 671 | 432 |

**Table 6.**
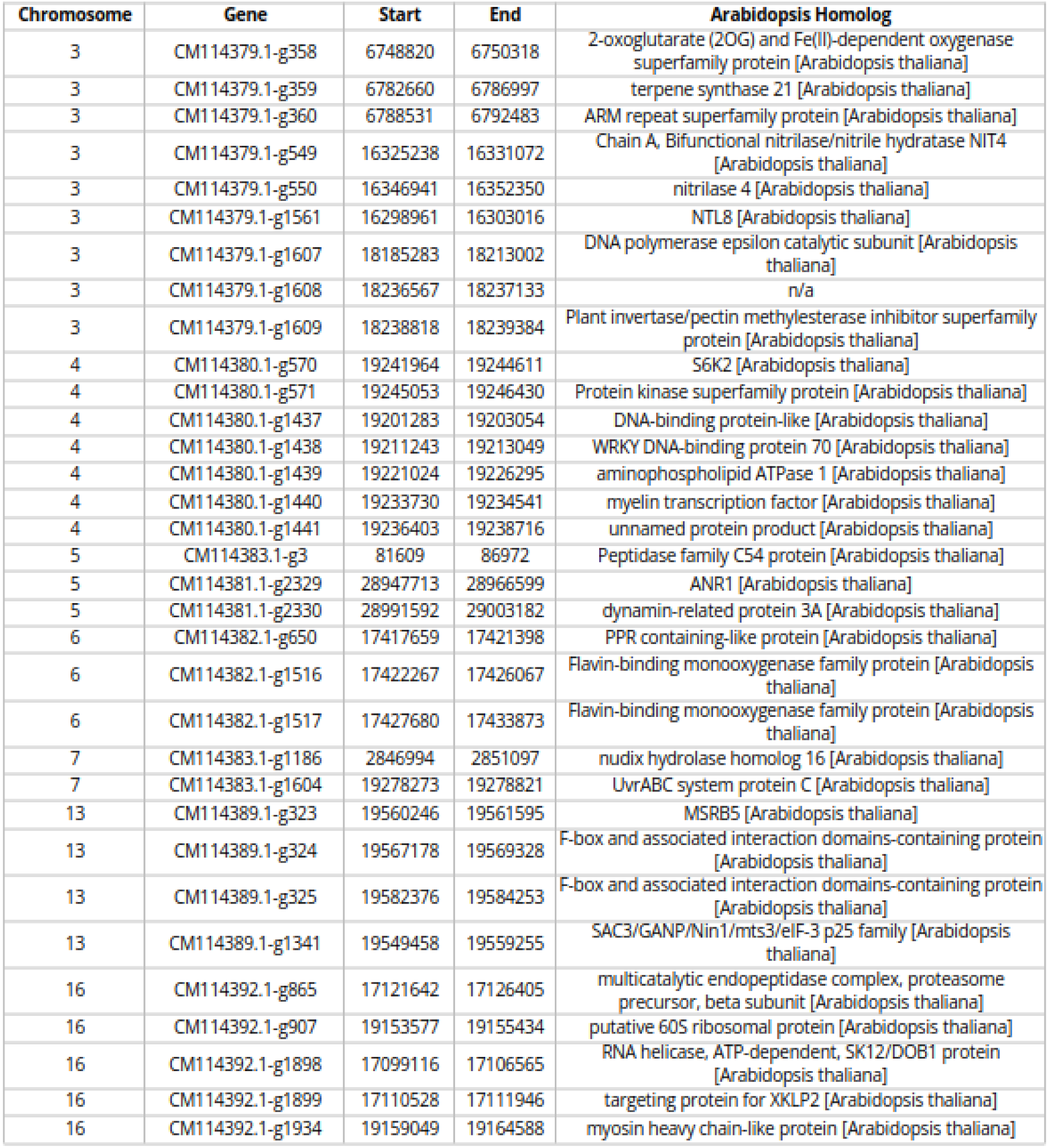
Genes in the Arid-Adapted *Pyrus* subgroup which are within the putative selective sweep regions, annotated with the closest *Arabidopsis* homolog.

## Conclusion

We developed a genus-scale pangenome for *Pyrus*, among the first such analyses in a tree fruit genus, in order to explore the origins of climate adaptation in *Pyrus*. Using this resource, regions of heightened diversity within the pangenome and evidence of selection could be examined for genes of interest in explaining *Pyrus* adaptation to diverse environments. This study also validates the viability of a single-platform sequencing approach in the large-scale *de novo* genomic assembly and analysis of land plant genera. This work provides a resource for future resequencing analysis for the genetic characterization of *Pyrus,* allowing for the alignment of sequencing reads with greatly reduced potential for reference bias. Augmentation of the pangenome graph with further genomes, transcriptomic data, and mapped resequencing data will be necessary to reach its full potential. Such a resource will enable connections to be made between genetic variants and phenotypes with genetic background context, which can advance the understanding of *Pyrus* genetics and enable efficient genetic improvement of cultivated *Pyrus* species.

## Author Contributions

June Labbancz: Conceptualization, Methodology, Investigation, Formal Analysis, Writing-Original Draft. Amit Dhingra: Conceptualization, Resources, Writing-Review and Editing, Supervision, Project Administration, Funding acquisition.

## Acknowledgements

The authors would like to thank the National Clonal Germplasm Repository (Corvallis, OR) staff for their help in accessing all the germplasm for this research. The authors are grateful to Professor Kate Evans at Washington State University for her critical reading of the draft of this manuscript.

